# Prolonged quiescence delays somatic stem cell-like division in *Caenorhabditis elegans* and is controlled by insulin signalling

**DOI:** 10.1101/462713

**Authors:** María Olmedo, Alejandro Mata-Cabana, María Jesús Rodríguez-Palero, Sabas García-Sánchez, Antonio Fernández-Yañez, Martha Merrow, Marta Artal-Sanz

## Abstract

Cells can enter quiescence in adverse conditions and resume proliferation when the environment becomes favourable. Prolonged quiescence comes with a cost, reducing proliferation potential and survival. Interestingly, cellular quiescence also occurs in normal development, with many cells spending most of their lifetime at this state. Elucidating the mechanisms involved in surviving long-term quiescence and in maintenance of cellular proliferation potential will contribute to a better understanding of the process of tissue regeneration. Developmental arrest of *C. elegans* at the L1 stage is an emerging model for the study of cellular quiescence and reactivation. During arrest, L1 larvae undergo a process that shares phenotypic hallmarks with the ageing of the adult. Interestingly, insulin signalling, a prominent pathway in the regulation of ageing, also balances cell proliferation and activation of stress resistance pathways during quiescence, becoming a candidate regulator of proliferation potential. Here we report that prolonged L1 quiescence delays reactivation of blast cell divisions in *C. elegans*, leading to a delay in the initiation of postembryonic development. This delay is accompanied by increased inter-individual variability. We propose that the delay in cell division results from the decline that animals suffer during L1 arrest. To that end, we show that insulin signalling modulates the rate of L1 ageing, affecting proliferative potential after quiescence. These findings support that the insulin signalling pathway has a comparable role in L1 arrest to that in ageing adults. Furthermore, we show that variable yolk provisioning to the embryos as a consequence of maternal age is one of the sources of inter-individual variability in recovery after quiescence of genetically identical animals. Taken together, these results support the relevance of L1 arrest as a model to study *in vivo* proliferation after quiescence and to understand the mechanisms for maintenance of proliferation potential.

## Introduction

In natural conditions, organisms are often subjected to changes in nutrient availability that modulate growth and proliferation. In the absence of growth-sustaining factors, cells can enter quiescence, a reversible, non-proliferative state. Under certain conditions, quiescent cells re-enter the cell cycle and resume proliferation ^1,2^. Despite the lack of proliferation, quiescence is not a homogeneous state. Cells move progressively into a deeper level of quiescence over time ^3^. As a consequence, the duration of quiescence affects cell viability and proliferation potential. Understanding the molecular processes that regulate the maintenance of proliferation potential and the mechanisms mediating reactivation of proliferation after quiescence is key in the comprehension of stem cell ageing. Furthermore, failure of the programs that negatively regulate cell division upon growth signal withdrawal can lead to uncontrolled proliferation and is a hallmark of cancer cells (reviewed in ^4^).

In the nematode *Caenorhabditis elegans*, newly hatched larvae enter a quiescent state when subjected to food deprivation. These L1 larvae (first larval stage) have 558 cells, 53 of which are somatic stem cells that divide further to reach the 959 somatic cells of the adult ^5^. These blast cells divide during four stages of postembryonic development (L1-L4), and somatic cell divisions end with the transition to adulthood. Arrest at the L1 stage involves quiescence of the blast cells that would normally divide during the first larval stage. The success of the arrest depends on the cyclin-dependent kinase inhibitor CKI-1/CIP/KIP/p27, that holds the cell cycle at G1 ^6^. The activation of CKI-1 in the absence of food is mediated by DAF-16/FOXO, a transcription factor activated in conditions of low insulin signalling ^7^. Arrested L1 larvae can survive several weeks without food and show increased resistance to stress ^8^. During L1 arrest, larvae undergo a process of ageing, manifested by the accumulation of protein aggregates, increased Reactive Oxygen Species (ROS) accumulation, and mitochondrial fragmentation. The addition of food to arrested L1 leads to the reversion of these ageing phenotypes and the initiation of larval development. However, extended L1 arrest reduces the potential to recover once food is available ^9^. Interestingly, the duration of L1 arrest has been proposed to affect the subsequent larval development. Following feeding after prolonged L1 starvation, animals take longer to reach adulthood and show increased variability in the time it takes to reach this developmental stage ^10,11^. For these reasons, the nematode offers an exceptional tractable model system to study the mechanisms that impact cell arrest and proliferation in a multicellular organism. Despite this interest, the processes taking place during L1 quiescence and recovery are poorly understood.

Here we show that extended L1 arrest delays the reactivation of the cell divisions marking the initiation of the postembryonic developmental program. Once the program is initiated developmental timing is not affected. This contrasts with previous interpretations concerning the delay in reaching adulthood observed in animals after prolonged starvation. Importantly, this finding suggests that prolonged starvation can lead to deeper quiescence of blast cells in *C. elegans*. In our effort to understand the signalling pathways controlling the depth of quiescence, we investigated the role of insulin signalling in recovering from L1 arrest. Low insulin signalling in the *daf-2* mutant leads to faster recovery and reduced ageing, probably indicating a shallower level of quiescence. Finally, we have discovered a source of variability in L1 quiescence by analysing the contribution of age-dependent maternal provisioning to the embryo. Larvae from older mothers, which provide more yolk proteins to the embryo, accumulate fewer ageing markers and recover faster from prolonged quiescence.

## Results and discussion

### Time to recover from L1 arrest increases with prolonged starvation and reflects proliferative potential after quiescence

When L1 larvae are arrested for prolonged periods of time, animals take longer to reach adulthood once they are fed ^10,11^. However, the duration of each of the four larval stages after extended starvation remains unexplored, hindering the analysis of the effects of starvation on the reactivation of the larval developmental program. Using a highly quantitative and novel assay for developmental timing ^12^, we measured the duration of all larval stages after different periods in L1 arrest (Fig. 1a). As previously observed, the entry into adulthood was increasingly delayed with longer times in arrest (Fig. 1b). When we analysed the duration of each stage of development individually, we found that all larval stages, except L1, and all molts were essentially unaffected by the duration of the arrest (Supplementary Fig. 1a). Instead, the delayed adulthood results from an extended recovery time, defined as the time from the exposure to food until the entry into the first molt (L1; Fig. 1c). Prolonged starvation also resulted in increased variance of recovery time (F test p-value <0.0001, when comparing day 2 and day 8) (Fig. 1c). Starvation time had no effect on the time between the first and the last molt (M1-M4) (Fig. 1d). In summary, the growth rate of the nematode is not affected by starvation time. Developmental timing is largely resilient to extended arrest, whereas recovery time increases with the duration of the arrest.

**Figure 1.**
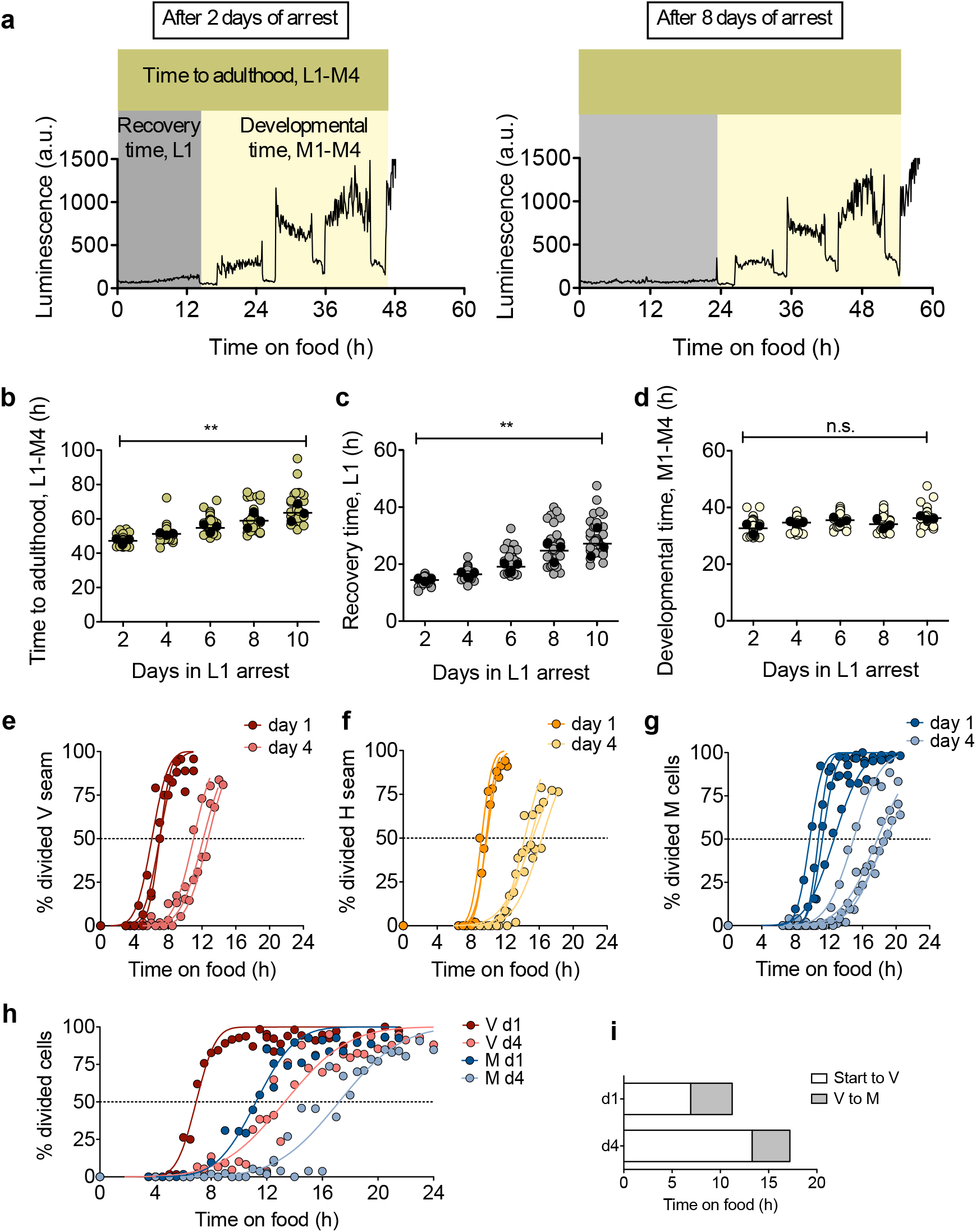
Prolonged quiescence delays reactivation of the developmental program. **a.** Representative plots of the duration of development for animals arrested as for 2 days (left panel) and 8 days (right panel). The different developmental stages quantified during the experiment are indicated. Recovery time (L1) is defined as the time between the addition of food to starved L1 animals and the initiation of the first molt. Developmental timing is defined as the period between the beginning of the first molt and the end of the last molt, or initiation of adulthood (M1-M4). Total time to reach adulthood is the time between the addition of food and the end of the last molt (L1-M4). **b-d.** Total time to adulthood (b), recovery time (c), and developmental time (d) for larvae arrested for 2, 4, 6, 8, or 10 days. Average values per experiment are indicated with a black dot and values from single animals are indicated with a coloured dot. We performed One-way ANOVA on the averages of 3 biological replicates (** p<0.01). The total number of animals is 47, 41, 45, 38, and 27 for samples from 2 to 10 days of arrest, respectively. **e-g.** Timing of seam/M cell division upon addition of food to L1 larvae arrested for one day or for four days. We analysed specific reporters for the division of seam cells from the V lineage (e), seam cells from the H lineage (f) and M cells (g). The plots show data from 3-4 biological replicates and values represent the percentage of animals showing division from a population of at least 40 larvae. Curves represent the fit assuming a cumulative Gaussian distribution. The dashed lines indicate the value of 50% of animals with divided cells. **h.** Timing of division of V seam cells and M cells upon addition of food to L1 larvae arrested for one day or for four days. We analysed a double reporter to assay both divisions in the same animals. The plots show data from two biological replicates and values represent the percentage of animals showing division from a population of at least 40 larvae. **i.** Prolonged arrest delays the division of V seam cells but not the timing between V and M cell divisions. The graph shows the time necessary to find 50% of animals with the corresponding divisions, estimated as the average of the cumulative Gaussian distribution of the data in panel h.

In order to understand the nature of the extended L1 stage, we fitted the recovery times after different starvation durations using linear regression. The stage L1, by definition, starts with hatching of the embryos and ends with the transition to L2. However, this definition does not consider the time needed to launch the postembryonic developmental program. As a consequence, L1 appears longer than the rest of larval stages. The relative molt/larva duration is markedly smaller for M1/L1, compared to all the other larval stages ^12^. From the linear regression, we calculated the time (*y* value) needed to reach the first molt when time in arrest (*x*) equals 0. This provides an estimate of the *bonafide* duration of L1 for non-starved larvae of *ca.* 10 hours (Supplementary Fig. 1B). With the *bonafide* L1 duration that we determined from our data, we could re-calculate the relative molt/larva duration and found out that, in this case, the M1/L1 ratio reaches the same value as all the other larval stages (Supplementary Fig. 1c). Our hypothesis is that prolonged L1 arrest does not affect the duration of the *bonafide* L1, but it determines the delay that occurs between re-introduction of food and reactivation of postembryonic cell divisions. However, an alternative explanation would be that, after extended L1 arrest, the reactivation of cell divisions would take place after the addition of food without a delay but then the complete L1 stage would be prolonged (Supplementary Fig. 1e). To differentiate between these two options, we analysed the relative timing of specific events during L1, namely, the divisions of seam and M cells. Seam cells undergo a single round of asymmetric self-renewing division at the beginning of each larval stage (L1-L4), immediately after ecdysis of the cuticle. The first of these divisions takes place about five hours after hatching. M blast cells present in the L1 larvae give rise to the complete mesodermal cell lineage. The first division of the M cell takes place midway through this larval stage ^5^. We monitored the timing of seam cells (both V and H lineages) and M cell division after one and four days of arrest, using reporter strains expressing *Pscm::gfp* and *Phlh-8::gfp*, respectively. After a single day of arrest, seam cells of the V1-4 and H lineages start to divide after five or eight hours, respectively, following exposure to food (Fig. 1e, 1f). M cells divide after nine hours (Fig. 1g). After four days of arrest, the divisions were delayed on average by 5.75 hours in the case of seam cells and 6.43 hours in the case of the M cells. These results already support the first scenario, in which relative timing of divisions is similar in animals arrested only briefly or for extended periods.

In order to be able to compare the time of these divisions within the same animals, we made a double seam and M cell reporter and monitored V and M cell divisions upon feeding of larvae arrested for one and four days (Fig. 1h). We then compared the time needed for 50% of animals to achieve these cell divisions in the two conditions. While the time between the addition of food and V cell divisions is almost doubled in L1s arrested for four days, the time between V and M cell divisions remains constant (Fig. 1i). We concluded that reactivation of blast cells is delayed after prolonged quiescence in *C. elegans*. This supports our hypothesis that the *bonafide* L1 is constant over time in arrest, as are the rest of larval stages. As a consequence, we can use recovery time to quantify proliferative potential after quiescence.

We asked whether this delay in cell divisions after prolonged arrest responded to a failure to release the repression exerted by DAF-16 in animals starved for four days. We observed that the number of animals with cytoplasmic localization increases rapidly in response to food independently of the duration of the arrest (Supplementary Fig. 1d). These findings confirm our hypothesis that exposure to food is not sufficient to resume the larval developmental program, instead reactivation of cell divisions is delayed after prolonged L1 arrest (Supplementary Fig. 1e), possibly reflecting a progressively deeper level of quiescence in these conditions.

### Insulin signalling modulates L1 ageing and recovery

One explanation for the delayed reactivation of cell divisions is that the arrested larvae need to counteract the damage caused during prolonged arrest. Since quiescent L1 develop age related phenotypes ^9^, we decided to investigate the process of L1 ageing and whether it influences reactivation of larval cell divisions. Insulin signalling is a prominent pathway in the control of ageing in the adult ^13^ and a modulator of the survival of arrested L1s ^14^. Under conditions of nutrient availability, the binding of insulin and insulin-like peptides to its receptor protein, called DAF-2 in *C. elegans*, provokes a conformational change of the intracellular portion of the receptor that leads to its phosphorylation. The activation of the receptor activates a pathway that leads to the phosphorylation of AKT proteins. AKT-1/AKT-2 phosphorylate DAF-16/FOXO that is retained in the cytoplasm and remains transcriptionally inactive. Reduced insulin/IGF-1-like signalling leads to activation of DAF-16/FOXO as it favours its translocation to the nucleus ^15-18^. In the absence of food, the transcription factor DAF-16 mediates both quiescence of the blast cells by the activation of the cyclin-dependent kinase inhibitor CKI-1, and the activation of stress resistance and maintenance pathways ^7,8^. Similar to what is observed with adult lifespan, reduction-of-function mutations in the gene encoding the insulin/IGF-1 *C. elegans* receptor *daf-2* increase survival during L1 arrest and mutations in *daf-16* reduce it ^7,14^. We measured recovery time in *daf-16(mu86)* and *daf-2(e1370)* mutants. After one day of starvation, the recovery time of the *daf-16* mutant was similar to that of the *wild-type* strain. However, after four days of starvation, the *daf-16* mutant showed a remarkable delay in the recovery time. For some animals, recovery took more than 50 hours (Fig. 2a, left). There is also an increased variability in recovery time among *daf-16* animals starved for four days (F test p-value <0.0001, when comparing *daf-16* to *wt*). Despite the long recovery time, the developmental timing of *daf-16* is only mildly delayed compared to that of the wild-type strain (Fig. 2a, right). This result suggests that DAF-16 contributes to maintain proliferation reactivation potential.

**Figure 2.**
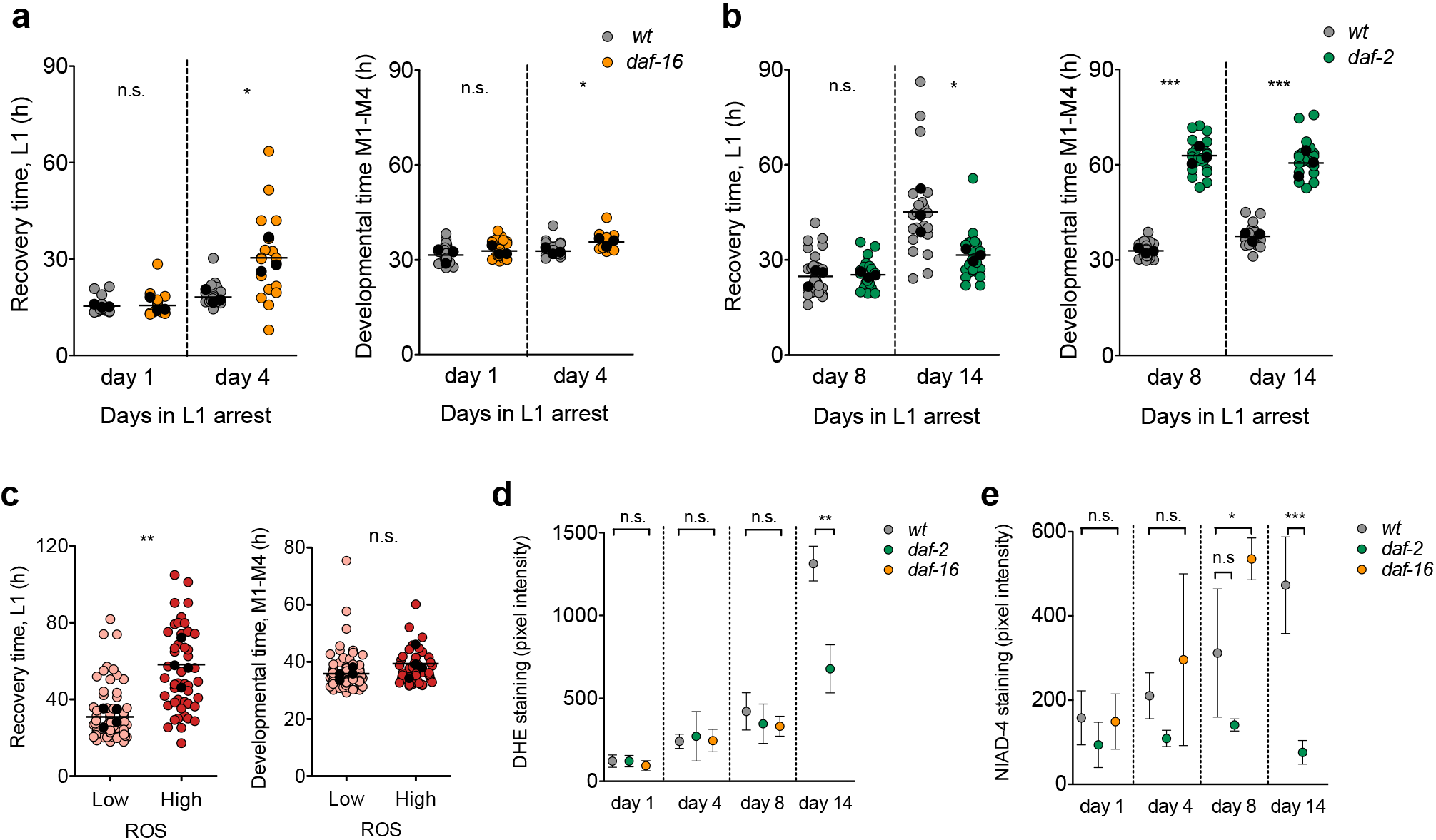
Low insulin signalling during quiescence ameliorates proliferative potential and attenuates L1 ageing. **a.** Recovery and developmental time for the wild-type strain and the *daf-16(mu86)* mutant after one and four days of arrest. Average values per experiment are indicated with a black dot, and values from single animals are indicated with a coloured dot. We performed t-test on the averages of 3 biological replicates (* p<0.05). The total number of animals, in the order shown in the plot, is 33, 27, 25 and 18. **b.** Recovery and developmental timing for the wild-type strain and the *daf-2(e1370)* mutant after 8 and 14 days of arrest. Average values per experiment are indicated with a black dot, and values from single animals are indicated with a coloured dot. We performed One-way ANOVA on the averages of 3 biological replicates (* p<0.05). The total number of animals, in the order shown in the plot, is 32, 24, 22 and 24. **c.** Recovery and developmental timing for animals categorized as having low or high DHE staining after 8 days of starvation. Average values per experiment are indicated with a black dot, and values from single animals are indicated with a coloured dot. We performed One-way ANOVA on the averages of 4 biological replicates (** p<0.01). The total number of animals for low and high DHE is 82 and 46, respectively. **d.** Quantification of ROS accumulation in the wild-type strain, *daf-2 (e1370)*, and *daf-16(mu86)*. Plots show mean (± s.d.) of the biological replicates. We measured DHE signal in the head of at least 15 animals per conditions in 3-4 biological replicates. For days 1, 4, and 8 we performed One-way ANOVA on the averages of biological replicates, followed by Dunnett’s Multiple Comparison test to detect significant differences between the mutants and the wild type. For day 14 we performed t-test (** p<0.01) **e.** Quantification of amyloids in the wild-type strain, *daf-2(e1370),* and *daf-16(mu86)*. We measured NIAD-4 signal in the head of at least 15 animals per conditions in four biological replicates. Plots show mean (± s.d.) of the biological replicates. For days 1, 4, and 8 we performed One-way ANOVA on the averages of biological replicates, followed by Dunnett’s Multiple Comparison test to detect significant differences between the mutants and the wild type animals (* p<0.05). For day 14 we performed t-test (*** p<0.001).

We then investigated whether increased DAF-16 activation, as that found in *daf-2(e1370)* mutants, leads to enhanced reactivation of proliferation. Interestingly, *daf-2* mutants produce longer, better provisioned embryos which, after prolonged starvation, produce more progeny than the wild type ^19^. When we measured recovery after one and four days of arrest we did not observed a faster recovery of the *daf-2* mutant (Supplementary Fig. 2a). However, when we maintained L1 larvae in starvation for 14 days, the *daf-2* animals recovered more rapidly than the wild type (Fig. 2b, left). This result is remarkable since low insulin signalling is generally related to proliferative defects ^20^. In these experiments, developmental time (M1 trough M4) in *daf-2* mutants showed a delay when compared with the wild-type strain (Fig. 2b, right), as previously described ^12,21^. Importantly, the duration of arrest did not have a strong impact on growth rate, neither in the wild type nor in the *daf-2* mutant (Fig. 2b, right). These results indicate that activation of DAF-16 confers an advantage for recovery, possibly by reducing the rate of L1 ageing. Despite the proliferative role of insulin signalling, its repression could be beneficial for maintenance of cells at a shallower level of quiescence, allowing faster recovery.

### Markers of ageing correlate with increased recovery time

In light of the previous results, we asked whether recovery time, as a proxy for reactivation of quiescent cells, correlated with the process of L1 ageing. We investigated if recovery time was related to the accumulation of ROS, one of the markers of L1 ageing ^9^. We stained L1 arrested larvae with Dihydroethidium (DHE), a compound that fluoresces in the presence of ROS. We observed an increasing variability in DHE signal over time of arrest (Supplementary Fig. 2b). After 8 days of L1 arrest, we could visually establish two categories of larvae with high and low fluorescence in the head, which yielded significant differences when we quantified the actual fluorescent signal (Supplementary figure 2c). We selected animals based on these two categories and analysed their recovery, demonstrating that animals with higher DHE signals recovered significantly more slowly than those with lower signals (Fig 2c. left) but did not show developmental delay (Fig. 2c, right). This result suggests that the capacity to recover depends on the accumulation of age-related phenotypes ^9^.

We suspected that insulin signalling modulates the rate of accumulation of L1 ageing markers. Therefore, we looked at the progression of ROS accumulation in the mutants *daf-2* and *daf-16* over the time in arrest. The increase in the DHE signal was less pronounced in the *daf-2* mutant, showing significant differences relative to the wild type animals after 14 days of arrest (Fig 2d). We were only able monitor DHE signals in the *daf-16* mutant up to day 8 of arrest, due to the increased mortality of these animals ^22^. At that time, the DHE signal was similar to that of wild type, suggesting that either ROS accumulation does not account for the delay in reactivation of proliferation of this mutant or DHE staining does not reflect all ROS present in the nematode. Next, we checked whether the delayed recovery of *daf-16* mutants is due to a pronounced accumulation of other markers of ageing. We analysed the formation of amyloids using the dye NIAD-4 ^23^. In this case, the differences in staining in the *daf-2* mutant after 14 days of arrest are even more pronounced. The mutant *daf-16* showed significant differences in NIAD-4 staining after 8 days of arrest (Fig. 2e).

Altogether, these results suggest that the physiological decline, similar to the ageing process, that takes place during L1 arrest leads to slower recovery once animals are fed. The decreased aging rate of *daf-2* mutants during L1 arrest possibly results from a combination of both a better provisioning from the mothers ^19^ and an optimized rationing of the available resources due to prolonged activation of DAF-16 targets during L1 arrest in insulin signalling mutants (Supplementary Fig. 2d and ^24^).

### Maternal age influences recovery time of its progeny

A recurrent observation for both recovery time and ageing markers is an increase in the coefficient of variation as the values increase. This means that processes that increase quiescence depth show an important interindividual variability. The causes of such variability in genetically identical organisms reared in the same environmental conditions are only starting to be elucidated. One factor introducing phenotypic variability in development after extended arrest is differential embryo provisioning of vitellogenin or yolk proteins, as a consequence of maternal age. Embryos from older mothers contain more yolk proteins and develop faster ^25^. We investigated whether this variable could lead to delays in reactivation of proliferation. We obtained embryos from gravid adults in their first, second, and third day of egg laying, maintained them in L1 arrest for 8 days and measured the development of individual larvae after providing them with food. Although variability was high in all conditions tested, we found that larvae from older mothers recovered from developmental arrest significantly faster than those of younger mothers (Fig 3a). On average the recovery times for larvae from mothers in the first, second and third days of egg production were 27.64, 22.40, and 21.70 hours respectively. Although larvae from day 3 mothers were on average similar to those from day 2 mothers, this population contained the fastest animals in terms of recovery. When analysing the 50% fastest animals in each condition, average recovery was progressively faster, with recovery times of 22.52, 19.65, and 17.56 hours respectively (Fig 3b). This effect was more prominent when we analysed recovery of the very first embryos produced by the mothers (day 0.5, Supplementary Fig. 3a). As in all cases we have seen, maternal age does not have an effect on developmental timing after extended arrest (Fig. 3c and Supplementary Fig. 3a).

**Figure 3.**
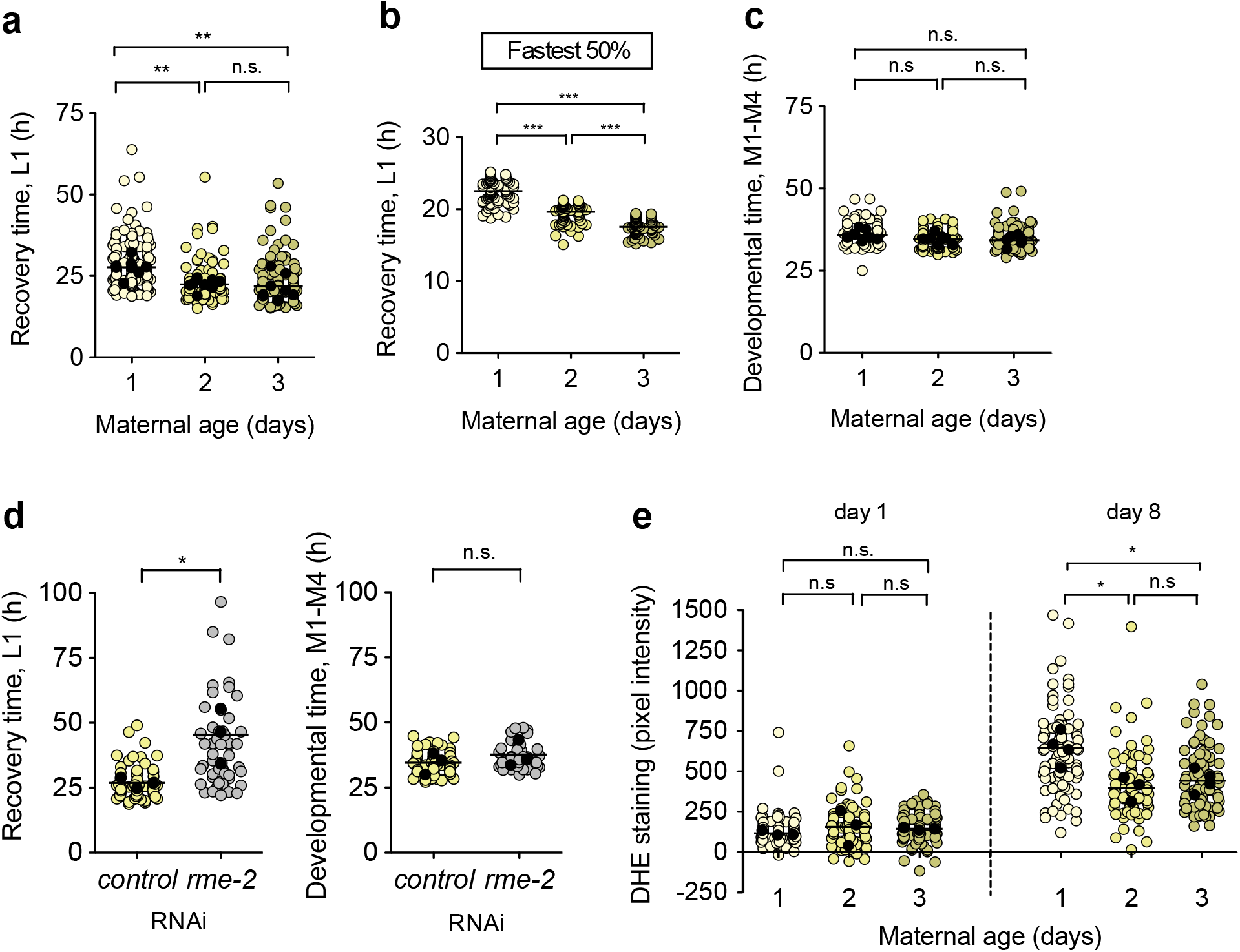
Maternal age introduces variability in recovery from larval arrest. **a.** Recovery time after 8 days of arrest of L1 larvae from day 1-3 progeny. Average values per experiment are indicated with a black dot, and values from single animals are indicated with a coloured dot. We performed One-way ANOVA followed by Bonferroni testing on the averages of 8 biological replicates (** p<0.01). The total number of animals is 159, 167 and 147 for samples from day 1 to 3. **b.** Recovery time of the fastest half of the population shown in (a). We performed One-way ANOVA followed by Bonferroni testing on the values from individual animals (*** p<0.001). **c.** Developmental time after 8 days of arrest of L1 larvae for day 1-3 progeny. Labelling and statistics are as in (a). **d.** Recovery time after 8 days of arrest for L1 larvae from mothers fed either control or *rme-2* RNAi bacteria. Average values per experiment are indicated with a black dot, and values from single animals are indicated with a coloured dot. We performed One-way ANOVA followed by Bonferroni testing on the averages of 3 biological replicates (* p<0.05). The total number of animals is 72 and 52. **e.** ROS accumulation after 1 or 8 days of arrest for day 1-3 progeny. We performed One-way ANOVA followed by Bonferroni testing on the averages of 3-4 biological replicates (* p<0.05). The total number of animals is 94, 84, 89, 102, 82 and 82.

We then investigated whether the reduction of vitellogenin/yolk proteins in the embryo affected recovery time after prolonged starvation. Yolk uptake by the oocyte occurs through RME-2 mediated endocytosis ^26^. Although yolk is not necessary for efficient reproduction in *C. elegans* ^27^, efficient yolk protein transport and storage contribute to L1 survival ^28^. We obtained L1 larvae after treatment of the mothers with *rme-2* RNAi and control bacteria and maintained them in L1 arrest for 8 days. When we measured development in response to food, we observed that larvae from embryos with reduced vitellogenin showed longer recovery times than larvae from mothers treated with control RNAi (Fig. 3d, left). Despite the pronounced delay in recovery, developmental timing was similar between conditions (Fig. 3d, right). Our results, therefore, confirm that yolk provisioning is important during L1 starvation but does not support a role in developmental timing *per se*.

We also analysed whether embryos from older mothers were protected against the accumulation of L1 ageing markers. We submitted L1 from day 1-3 mothers to arrest and measured DHE signals. We observed no significant differences after one day of arrest but a marked, significant difference after 8 days. Indeed, larvae from day 2-3 mothers showed reduced DHE accumulation compared to larvae from day 1 mothers (Fig 3e). We found no differences in amyloids, reported by NIAD-4 staining (Supplementary Fig. 3b). These results indicate a role of maternal age in preventing L1 ageing, leading to maintenance of proliferation potential. To explore a possible link between insulin signalling and yolk provisioning, we analysed the localization of DAF-16 during L1 arrest, in larvae treated with *rme-2* RNAi (Supplementary Fig. 3c). Larvae with reduced yolk provisioning had similar levels of nuclear DAF-16 than those treated with control RNAi. This result indicates that reduced L1 ageing of older mothers is unlikely to be due to an increased activation of DAF-16.

Altogether, our results reinforce the role of *C. elegans* as a model to study pathways involved in blast cell quiescence and advance our knowledge of this process, which is key to understanding the maintenance of proliferation potential in quiescent cells. The balance between cell quiescence and proliferation is crucial for maintenance of stem cell pools, which hold the potential to maintain tissue homeostasis or replace dead cells after injury. On the one hand, stimulation of proliferation of quiescent cells provokes the exhaustion of stem cells ^29^. On the other, the degenerative processes in stem cells and the systemic cues that regulate their activity have been connected to the age-dependent decline in regenerative potential of tissues ^30^. Proliferative potential of stem cells is reduced over time in quiescence. Now we have shown that, also in *C. elegans*, prolonged arrest delays blast cell divisions, suggesting that cells enter a deeper level of quiescence. Aged stem cells accumulate Reactive Oxygen Species (ROS), DNA damage, aggregated proteins and show mitochondrial misfunction (reviewed in ^31^). Interestingly, a recent work has determined that lysosomal gene expression increases as quiescence deepens. Lysosomal function favours a shallower level of quiescence by reducing ROS accumulation. Furthermore, a score for gene expression during the deepening of quiescence parallels cellular senescence and aging ^32^. This recent finding and our own results suggest that ROS accumulation favours deeper quiescence. Furthermore, DAF-16 drives lysosomal function in *C. elegans* ^33^ which could explain the observation that *daf-2(e1370)* mutants, which present a constitutive DAF-16 activation, show lower levels of ROS and faster recovery. Although FoxO deficient Hematopoyetic Stem Cells (HSC) present increased ROS ^34^, we do not observed higher ROS accumulation in a *daf-16* mutant. However, it is possible that other factors are limiting survival of this mutant before the increased accumulation of ROS becomes evident.

Roux et al. proposed that quiescent L1 suffer a physiological decline similar to the ageing of the adult. Most ageing markers accumulated by L1 larvae during arrest were reversed by feeding, and the competence of the animal to clear those signs of ageing determines its capacity to recover from L1 arrest. Here we show that we can measure the proliferative potential of quiescent L1, not only in terms of whether they recover or not, but precisely quantifying it as the recovery time. This way, we have been able to quantify the recovery of *daf-2* mutants, showing that this long-lived mutant has increased proliferation capacity after prolonged quiescence. DAF-16 maintains cellular arrest and favours survival through independent mechanisms; disruption of the pathways that allow blast cell division in the absence of DAF-16 does not enhance survival of the *daf-16* mutant ^7,22^. Now we show that activation of DAF-16 slows the accumulation of L1 ageing markers, probably promoting a shallower level of quiescence that permits faster recovery.

We have also found that larvae from older mothers present less ROS accumulation and recover faster from quiescence, suggesting that they are at a shallower level of quiescence. Whether ROS levels and yolk provisioning are mechanistically link requires further investigation. We did not find alterations in DAF-16 activation in yolk depleted larvae and *daf-2* mutants produce less yolk proteins^35,36^. These observations suggest that the mechanisms for ROS reduction in conditions of low insulin signalling and in older mothers likely have different origins.

In sum, our results support the relevance of L1 arrest upon starvation as a model to study proliferative potential after quiescence *in vivo*.

## Materials and methods

### Culture conditions and strains

We cultured stock animals according to standard methods ^37^, maintaining them at 20 °Con nematode growth medium (NGM) with a lawn of *Escherichia coli* OP50-1. The only exception is in Fig. 1a-d, where animals were maintained at 18 °C previous to the experiment. For the initial characterization of the recovery time and developmental timing in response to starvation (Fig. 1a-d), we used the reporter strains PE255 *feIs5* [*Psur-5::luc+::gfp; rol-6 (su1006)*]X ^38^. For subsequent experiments we created a new luciferase reporter without the marker *rol-6*, generating the strain MRS387 *sevIs1 [Psur-5::luc+::gfp]X*. To generate MRS387 we injected the plasmid pSLGCV in N2 worms and selected several lines that transmitted the array. We then irradiated one of these strains MRS222 *sevEx1[Psur-5::luc+::gfp]* with X-rays to integrate the reporter array. These strains were outcrossed 10 times with N2. We crossed the new reporter strain using standard genetic techniques to generate MRS424 *daf-16(mu86); sevIs1 [Psur-5::luc+::gfp]X* and MRS434 *daf-2(e1370); sevIs1 [Psur-5::luc+::gfp]X*.

For the analysis of cell divisions in response to feeding we used the strains GAL69 *matIs38 [Pscm::CYB-1 DB-mCherry::unc-54 3′ UTR; Pscm::NLS-GFP::tbb-2 3′ UTR; Pmyo-2::GFP]*, PD4666 *ayIs6 [hlh-8::GFP fusion + dpy-20(+)],* JR667 *unc-119(e2498::Tc1)III; wIs51[SCMp::GFP + unc-119(+)]V* and the double seam and M cell reporter MOL198 *matIs38 [Pscm::CYB-1 DB-mCherry::unc-54 3′ UTR; Pscm::NLS-GFP::tbb-2 3′ UTR; Pmyo-2::GFP]*; *ayIs6 [hlh-8::GFP fusion + dpy-20(+)].* For the analysis of ROS and accumulation of amyloids, we used the strains N2, CB1370 *daf-2(e1370)*, and CF1038 *daf-16(mu86)*. To assess the localization of DAF-16 we used TJ356 *zIs356[Pdaf-16::daf-16a/b-gfp; rol-6]* IV. As a control for yolk reduction by RNAi treatment, we used RT130 *unc-119(ed3); pwIs23 [Pvit-2::vit-2::gfp, unc119(+)].*

### Preparation of starved L1

For starvation experiments we first treated gravid adults with alkaline hypochlorite solution to obtain embryos. For all experiments, we adjusted the concentration of embryos to 20 per microliter of M9 buffer. The embryos were incubated at 20 °C, with gentle shaking, leading to hatching and arrest at the L1 stage. After the corresponding amounts of time, arrested L1 were used for the different experiments. For recovery experiments, we staggered the hypochlorite treatments to be able to analyse, simultaneously, animals arrested for different periods of time. For the analysis of ROS and protein aggregation during arrest we followed the same cohort over the relevant periods of time, usually monitoring at day 1, 4, 8, and 14 of L1 arrest.

### Luminometry of single worms

We measured recovery time and developmental timing using a bioluminescence-based method that we previously developed ^12^. Briefly, L1 arrested animals were placed individually in wells of a white 96-well plate containing 100 μl of S-basal (including 5 μg/ml cholesterol) with 200 μM Luciferin. After all animals were placed in the wells, we added 100 μl of S-basal containing 20 g/l *E. coli* OP50-1 per well to resume development simultaneously for all the animals. Each well therefore contained 200 μl of S-basal with 10 g/l *E. coli* OP50-1 and 100 μM Luciferin. We then sealed the plate with a gas-permeable cover (Breathe Easier, Diversified Biotech) and measured luminescence (Berthold Centro LB960 XS^3^) for 1 second, at 5-minute intervals. We performed all experiments inside temperature-controlled incubators (Panasonic MIR-154/MIR-254) such that samples were held at 20 °C. We measured the duration of each larval stage and molt analysing the data as described in ^12^. We used sample sizes >20 individually quantified animals in at least 3 biological replicates. We alternated the samples across the plate to avoid local effects (i.e. temperature of the reader)

### Analysis of seam and M cell divisions

For the experiments with the single reporters, we obtained about 300 μl of L1 larvae suspension of the strains GAL69, JR667 (seam cell reporters) or PD4666 (M cell reporter) arrested for one day or four days. On the day of the experiment, we added the same volume of 20 g/l *E. coli* OP50-1 in S-basal (with cholesterol) to the arrested L1, to have a final concentration of 10 g/l *E. coli* OP50-1. At the indicated times, we collected 30 μl to a clean 1.5 ml tube and centrifuged for 1 minute at 3,000 r.p.m., in a table-top centrifuge. We removed 25 μl of supernatant and added 1 μl of 100 mM Levamisole. After the addition of food, cell divisions were monitored every hour over 3-14.5 hours (V seam cells) or 6.5-20.5 hours (H seam cells and M cells). Sampling times were displaced in time among independent experiments. We used this strategy to maximize the information to be obtained from the time course, since we are interested in distribution over time more than in each individual value. We calculated the percentage of animals with divisions over a population of at least 40 animals per time point and condition.

For the experiments with the double seam and M cell reporter, we prepared embryos twice per condition, separated by 12 hours. This way we could resume development of these animals with a 12-hour difference, allowing monitor divisions in the first 24 hours over an experimental period of 12 hours. We obtained L1 larvae of the strain MOL198 arrested for one day or four days and then resumed development by the addition of food, as indicated above. We calculated the percentage of animals with divisions over a population from at least 40 animals per time point and condition. In order to calculate the time at which 50% of the developing population had divisions, we fitted the data to a cumulative Gaussian distribution and calculated the mean value. In both cases we monitored the animals using a Leica scope DMi8 using GFP excitation/emission filters.

### Subcellular localization of DAF-16

We obtained L1 larvae of the strain TJ356 arrested for one day or four days. As for the monitoring of cell divisions, we added one volume of 20 g/l *E. coli* OP50-1 in S-basal (with cholesterol) to the arrested L1, to have a final concentration of 10 g/l *E. coli* OP50-1. At the indicated times, we took an 18 μl-aliquot of the suspension and placed it on a microscope slide with 2 μl of 10 mM Levamisole. We used a reduced concentration of Levamisole in order to avoid artificial translocation of DAF-16. We visualized the localization of DAF-16 in a Leica scope DMi8 using GFP excitation/emission filters and counted the animals with nuclear, cytoplasmic or intermediate localization. Since the localization of DAF-16 can be affected by temperature, the temperature of the room was maintained at 20 °C during visualization of the animals. We categorized animals as having nuclear, intermediate or cytoplasmic localization, in a population of at least 100 larvae. Since this categorization is somehow subjective, the experimenter was blind to the condition tested.

### DHE staining and longitudinal analysis of recovery

We added Dihydroethidium (DHE, Sigma) at a final concentration of 10 μM in 30 μl of arrested L1. After incubation for 2 hours at 20 °C, we centrifuged the sample, removed 25 μl of supernatant and added 1 μl of 100 mM Levamisole. We placed 5 μl of the anesthetized L1’s on a microscope slide with a cover slip. We imaged the animals in a Leica scope DMi8 at 200x magnification using transmitted light and mCherry excitation/emission filters. For quantification, we drew ROIs around the head of the larvae using the image from transmitted light and then quantified pixel intensity in the red channel using Image J. We subtracted the background for each image. We measured DHE of at least 15 animals per conditions in 3-4 biological replicates.

For the longitudinal analysis of recovery, we stained L1 larvae arrested for 8 days with DHE, as above, and then categorized them into animals with low or high signal. We placed these animals in individual wells of a 96 well plate containing 100 μl of S-basal with 200 μM Luciferin. After all animals were placed in wells, we added 100 μl of S-basal containing 20 g/l *E. coli* OP50 and proceeded as normal for luminometry of single animals. As above, during the experiment, wells contained 200 μl of S-basal with 10 g/l *E. coli* OP50-1 and 100 μM Luciferin. The plate was sealed, placed in the luminometer and processed as previously indicated.

### NIAD-4

We added NIAD-4 (Cayman Chemicals) at a final concentration of 1 μM (0.1% DMSO in M9 buffer) to an aliquot of 30 μl of arrested L1. After incubation for 2 hours at 20 °C, we centrifuged the sample, removed 25 μl of supernatant, and washed with 25 μl of M9 buffer to remove the excess NIAD-4. After centrifugation, we removed again 25 μl and added 1 μl of 100 mM Levamisole. We placed 5 μl of the anesthetized L1 on a microscope slide for analysis. We imaged the animals in a Leica scope DMi8 at 200x magnification using transmitted light and mCherry excitation/emission filters. Quantification was performed as for DHE staining. We measured DHE of at least 15 animals per conditions in four biological replicates.

### Synchronization of mothers

To obtain embryos from mothers in their first, second and third day of egg laying we prepared synchronized populations allowing 20 gravid adults to lay eggs on NGM plates for two hours. After this period, we removed the gravid adults and allowed the progeny to grow at 20 °C. After approximately 80, 88, 104, 112, 128, and 136 hours the progeny were respectively at what we have named day 0.5, 1, 1.5, 2, 2,5 and 3 of egg laying. Day 0.5 refers to the time when the first embryos have been produced, while at day 1 all animals (initially synchronised by 2 hours) have started egg laying. We staggered this process of obtaining synchronized mothers in order to perform the hypochlorite treatments of the three conditions on the same day.

### Yolk reduction by *rme-2* RNAi

We obtained embryos with reduced yolk by alkaline hypochlorite treatment of gravid adults grown on *rme-2* RNAi from the Ahringer library. We cultivated *rme-2* RNAi bacteria overnight and diluted 1/10 with control bacteria HT115 containing the empty plasmid pL4440. We added 500 μl of the diluted RNAi or control bacteria on NGM plates with 1 mM IPTG and 100 μg/ml ampicillin. We let the bacterial lawn dry and incubated the plates for 5 hours at 37 °C and overnight at room temperature. We transferred 20 gravid adults of the strain MRS387 per plate and let them lay eggs for 2 hours before retiring them from the plates. We grew them for 4 days at 20 °C before proceeding with the hypochlorite treatment. We maintained the animals in L1 arrest for 8 days at 20 °C. Development was re-initiated by adding OP50-1 bacteria and monitor development as previously described. As a control for the RNAi treatment we used the strain RT130 to monitor VIT-2::GFP defective transport into oocytes.

### Data analysis and statistics

We analysed luminometry data as previously described ^12^. For all luminometry experiments we have plotted the values for each individual animal and also the average of the values for each biological replicate. For statistics we have used the averages of independent biological replicates to avoid the inflated N value from using individual animals. The averages of most groups were normally distributed but in cases when N was too low to assess normality we visually inspected the values to discard important deviations from normality and variances were similar. We used unpaired two-tailed t-test to compare the means of two groups and one-way ANOVA to compare more than two groups. ANOVA was followed by Bonferroni’s post-hoc to compare all groups or by Dunnett’s test to compare groups to a control condition (mutants vs. wild type). For the analysis of DAF-16 localization in wild type and *daf-2* mutants over time in arrest we have performed Two-way ANOVA.

## Acknowledgements

Some strains were provided by the *Caenorhabditis* Genetics Center (CGC), which is funded by NIH Office of Research Infrastructure Programs (P40 OD010440). The strain PE255 was a gift from Cristina Lagido (University of Aberdeen) and GAL69 was a gift from Matilde Galli (Hubrecht Institute, Utrecht). M.A-S and M.O are supported by the Ramón y Cajal program of the Spanish Ministerio de Economía y Competitividad, RYC-2010-06167 and RYC-2014-15551, respectively. Our work is supported by the Spanish Ministerio de Economía y Competitividad (BFU2016-74949-P and BFU2012-35509), the European Research Council (ERC-2011-StG-281691), and a Marie-Curie Intra-European Fellowship (FP7-PEOPLE-2013-IEF/GA Nr: 627263).

## Author contributions

M.O., A.M-C, M-J.R-P, S. G-S, and A. F-Y carried out the experiments; M.O. and A.M-C designed the experiments; M.O, A.M-C, M-J.R-P, M.M and M.A-S interpreted results; and M.O, A.M-C, M.M and M.A-S wrote the manuscript.

---

**Figure S1. Relative to figure 1.**
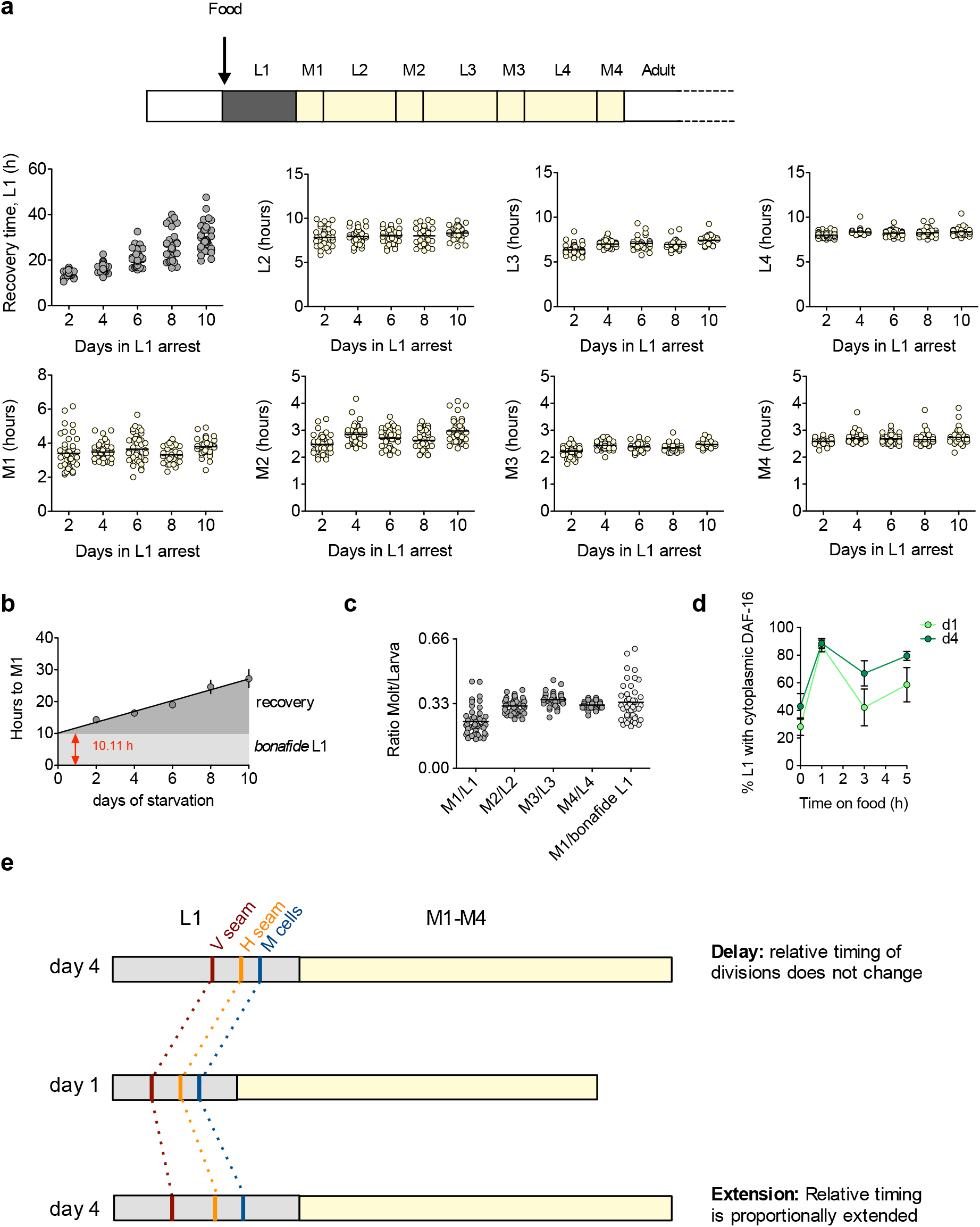
**a.** Duration of each stage of development upon feeding after 2-10 days of L1 arrest. Larval stages (L1-L4) and molts (M1-M4). **b.** Linear regression of the average values in Fig. 1c and estimation of the bonafide L1 as the time necessary to enter the first molt from the point of arrest. **c.** Molt/larva ratios for all larval stages using the actual data after two days of arrest or using the calculated *bonafide* L1. **d.** Percentage of animals with cytoplasmic localization of DAF-16 in the first five hour upon the addition of food after one or four day of L1 arrest. The plot shows the mean and SEM of four biological replicates. **e.** Alternative explanations of the longer recovery time (L1) after prolonged quiescence. Our data fits with a delay in the initiation of postembryonic development.

**Figure S2. Relative to figure 2.**
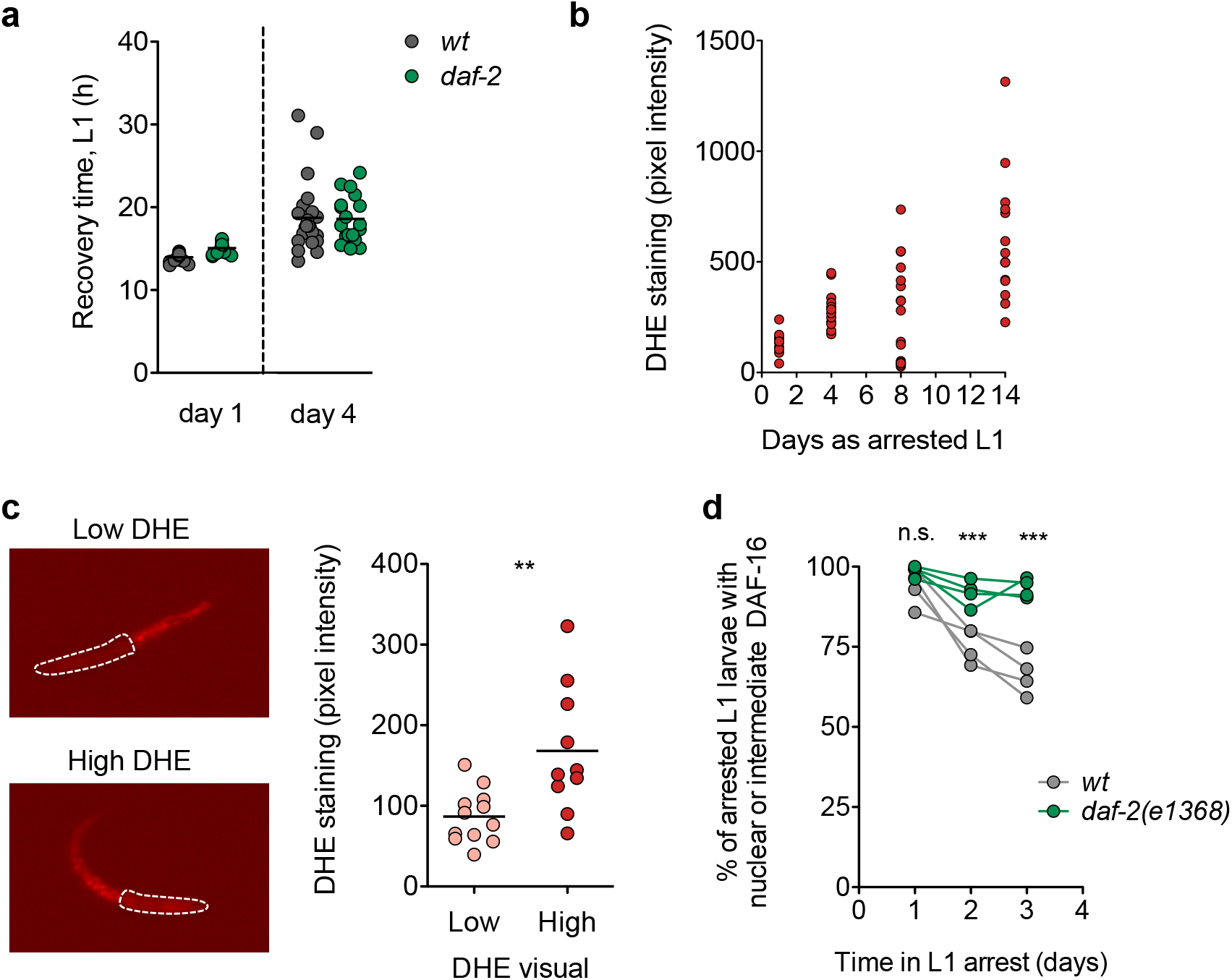
**a.** Recovery for the wild-type strain and the *daf-2(e1370)* mutant after one and four days of arrest. **b.** ROS accumulation over time in L1 arrest. The plot shows data from at least 10 animals per condition. **c.** Representative pictures of animals with low and high DHE signal after 8 days of starvation (left panel) and quantification of DHE staining in animals visually categorized as having low or high signal (right panel). **d.** Percentage of *wt* and *daf-2(e1368)* L1 larvae with nuclear or intermediate localization of DAF-16::GFP. We performed Two-way ANOVA followed by Bonferroni testing on the data from four biological replicates (*** p<0.001).

**Figure S3. Relative to figure 3.**
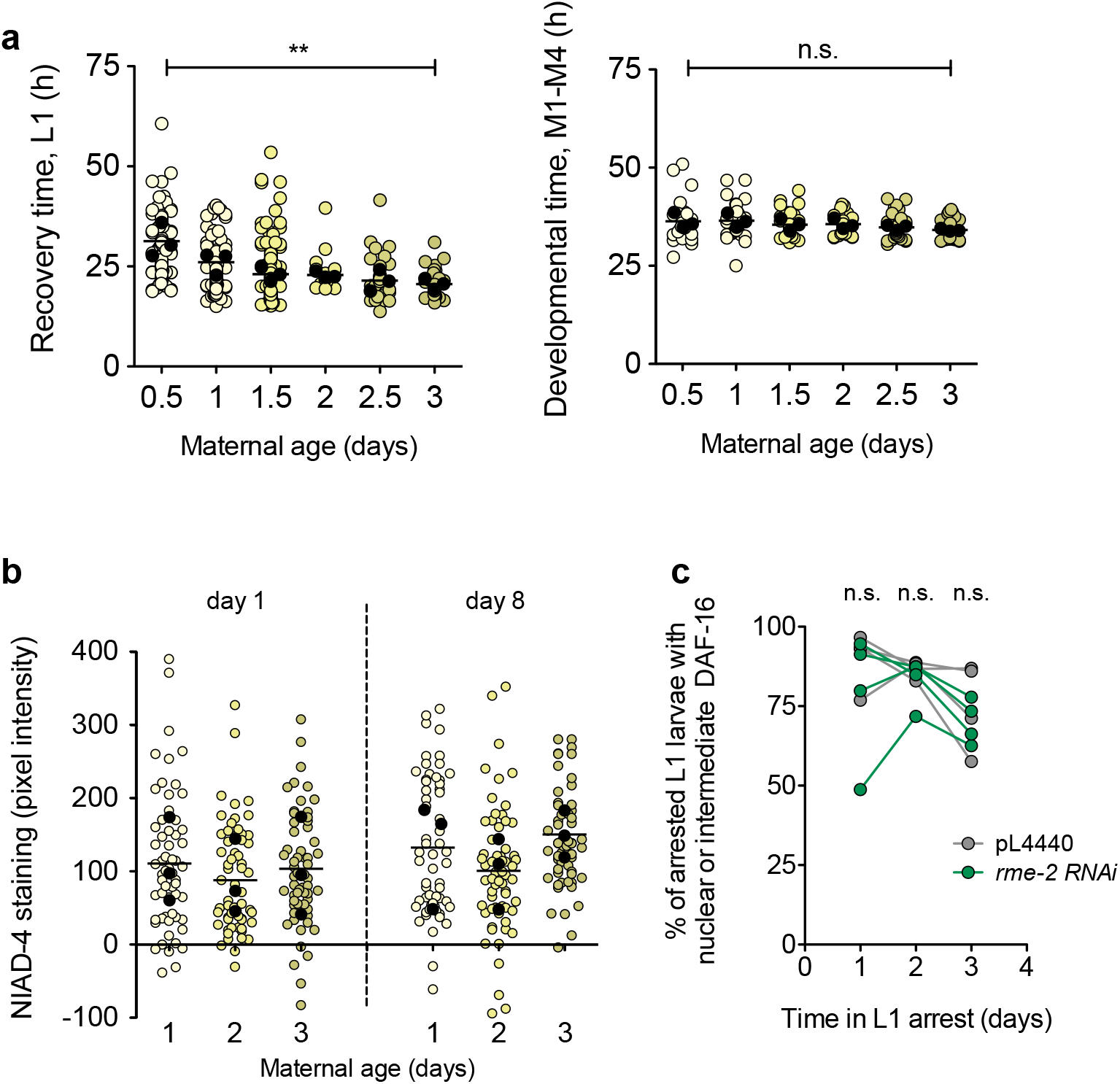
**a.** Recovery time and developmental timing after eight days of L1 arrest from progeny collected at the indicated maternal age (0.5-3 days of egg laying). Average values per experiment are indicated with a black dot, and values from single animals are indicated with coloured dots. We performed One-way ANOVA followed by Bonferroni testing on the averages of 3 biological replicates (** p<0.01). **b.** NIAD-4 accumulation in day 1-3 progeny after one and eight days of arrest. The plot shows data from 3 biological replicates. Average values per experiment are indicated with a black dot, and values from single animals are indicated with coloured dots. **c.** Percentage of L1 larvae with nuclear or intermediate localization of DAF-16::GFP, for animal treated with pL4440 or *rme-2* RNAi. We performed Two-way ANOVA followed by Bonferroni testing on the data from four biological replicates.

